# Strategies of tolerance reflected in two North American maple genomes

**DOI:** 10.1101/2021.07.19.452996

**Authors:** Susan L. McEvoy, U. Uzay Sezen, Alexander Trouern-Trend, Sean M. McMahon, Paul G. Schaberg, Jie Yang, Jill L. Wegrzyn, Nathan G. Swenson

**Author notes:** Corresponding authors: Jill Wegryzn, Nate Swenson.

## Abstract

Maples (the genus *Acer*) represent important and beloved forest, urban, and ornamental trees distributed throughout the Northern hemisphere. They exist in a diverse array of native ranges and distributions, across spectrums of tolerance or decline, and have varying levels of susceptibility to biotic and abiotic stress. Among *Acer* species, several stand out in their importance to economic interest. Here we report the first two chromosome-scale genomes for North American species, *Acer negundo* and *Acer saccharum*. Both assembled genomes contain scaffolds corresponding to 13 chromosomes, with *A. negundo* at a length of 442 Mb, N50 of 32 Mb and 30,491 genes, and *A. saccharum* at 626 Mb, N50 of 46 Mb, and 40,074 genes. No recent whole genome duplications were detected, though *A. saccharum* has local gene duplication and more recent bursts of transposable elements, as well as a large-scale translocation between two chromosomes. Genomic comparison revealed that *A. negundo* has a smaller genome with recent gene family evolution that is predominantly contracted and expansions that are potentially related to invasive tendencies and tolerance to abiotic stress. Examination of expression from RNA-Seq obtained from *A. saccharum* grown in long-term aluminum and calcium soil treatments at the Hubbard Brook Experimental Forest, provided insights into genes involved in aluminum stress response at the systemic level, as well as signs of compromised processes upon calcium deficiency, a condition contributing to maple decline.

**Significance statement:** The first chromosome-scale assemblies for North American members of the *Acer* genus, sugar maple (*Acer saccharum*) and boxelder (*Acer negundo*), as well as transcriptomic evaluation of abiotic stress response in *A. saccharum*. This integrated study describes in-depth aspects contributing to each species’ approach to tolerance and applies current knowledge in many areas of plant genome biology with *Acer* physiology to help convey the genomic complexities underlying tolerance in broadleaf tree species.

## Introduction

*Acer saccharum* (sugar maple) is a long-lived and dominant species in New England forests with a native range representing Eastern Canada, and the Northcentral and Northeastern United States. Widely known for its vibrant autumn hues, high quality timber, and as the preferred species for the production of maple syrup, *A. saccharum* also plays a key role in its native ecosystems, altering soil mineral content (Lucash et al., 2012), moisture levels (Emerman & Dawson, 1996), and mycorrhizae communities (Cong et al., 2015). *A. saccharum* provides food and shelter to many mammals (Godman et al., 1990), resident and migratory birds (Darveau et al., 1992), and over 300 species of caterpillar. Although seeds from this predominantly monoecious species are wind dispersed, the early flowers are an important pollen source for bees in late winter (Blitzer et al., 2016).

One of the most phylogenetically (∼50 MY) and morphologically distinctive *Acer* species from *A. saccharum* is *Acer negundo* (box elder). *A. negundo* is a short-lived tree that has the largest range of all North American *Acer*. It is native to predominantly lower elevation regions of Canada, the United States, and Mexico (Figure 1; Enquist et al., 2016; Maitner et al., 2018). This adaptable pioneer species is often seen in disturbed sites and urban settings. *A. negundo* has soft wood, less concentrated sugars for syrup, grows rapidly, and can tolerate low nutrient soils, moderate salinity, and drought conditions. Its invasive status in large portions of Europe, South America, and Asia are indicative of greater phenotypic plasticity (Lamarque et al., 2013). *A. negundo* has a number of distinctive attributes including dioecy (Renner et al., 2007) and pinnately compound leaves, mostly seen only in close relatives. Within such a significant genus, these two species reflect a breadth of social and ecological diversity and importance that recommends better understanding of their genetic distinctions.

**Figure 1.**
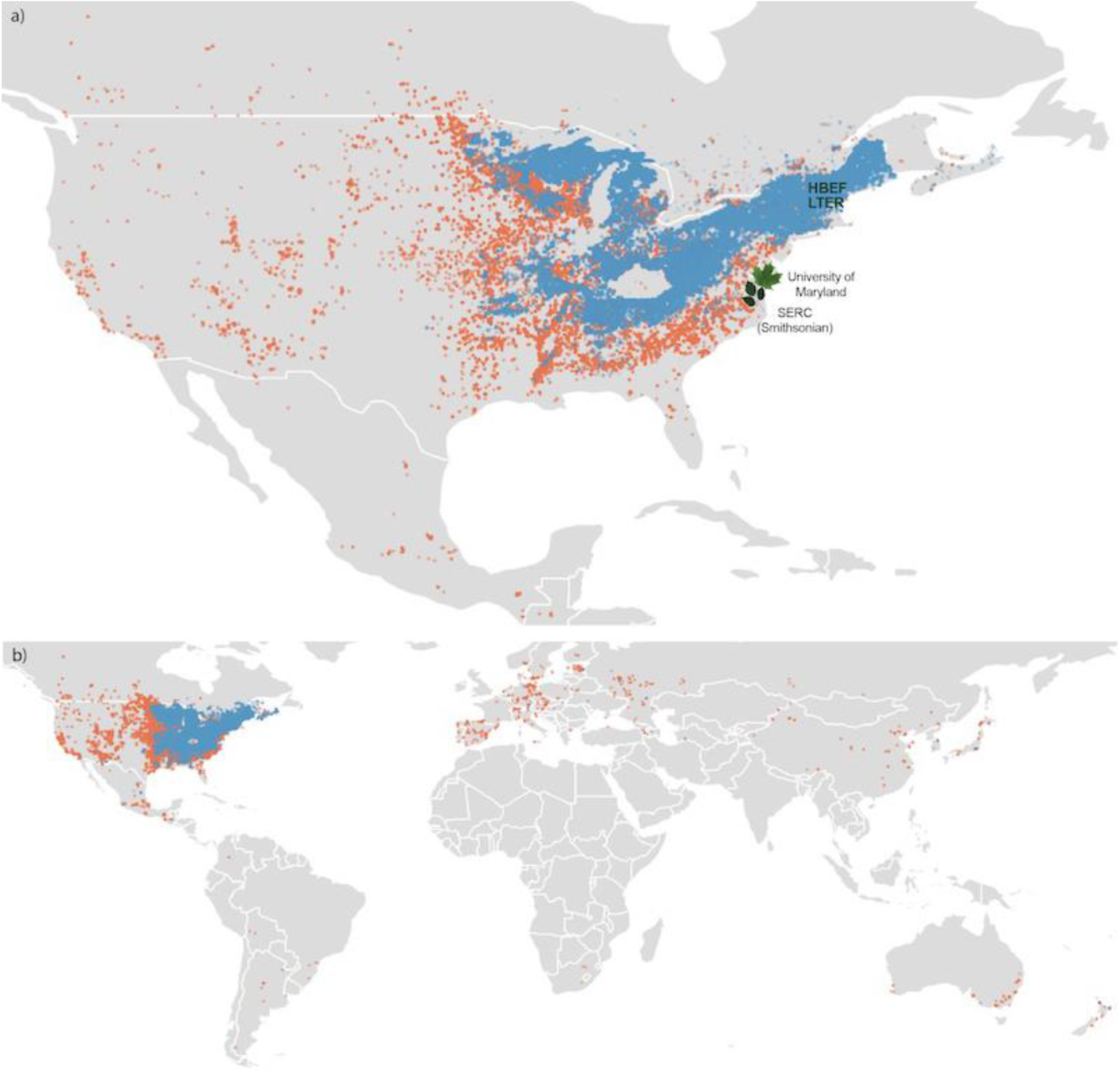
a) Native distributions of *A. saccharum* (blue) and *A. negundo* (orange) in North America. Leaves indicate location of individuals selected for the reference genomes; *A. saccharum* from the University of Maryland campus, and *A. negundo* from the Smithsonian Environmental Research Center. HBEF (Hubbard Brook Experimental Forest) is the location of the 9 individuals used for RNA-seq. b) All records of occurrence, native, introduced, and unknown, per BIEN 4.2. Non-native occurrences are predominantly *A. negundo*. https://gitlab.com/PlantGenomicsLab/AcerGenomes/-/blob/master/acer/supplemental/figures/acerranges.jpg

Forest ecosystems of the Northeastern U.S. are facing significant changes in composition driven by climate change (Rogers et al., 2017). “Maple decline” is a term referring to the loss of maple populations, originally referring to *A. saccharum*, but now applicable to *A. platanoides* and *A. rubrum* in the Northeast, and most recently, *A. macrophyllum* in the Northwest. Loss of *A. saccharum* has been documented over the last century, beginning in the late 1950s, leading to the first comprehensive, multidisciplinary study of this condition (Giese & Benjamin, 1964; Horsley et al., 2002). Maple decline is characterized by crown dieback, reduction in overall health and vigor, and a decrease in regeneration (Bishop et al., 2015). Episodic decline has increased in recent decades (Oswald et al., 2018). Decline and crown dieback of dominant *A. saccharum* provides a release for sympatric species such as *Fagus grandifolia* (American beech) which displays a higher level of tolerance to soil conditions and foliar aluminum ratios leading to shifting forest composition (Halman et al., 2015). Studies examining potential factors of maple decline have largely agreed that modified soil conditions, largely due to acid deposition, are the leading cause, compounded by additional climatic, pathogenic, and anthropogenic stressors (Bal et al., 2015). Acidic soils rapidly leach the essential cations calcium, magnesium and potassium, while mobilizing aluminum within the soil and contributing to more phytotoxic forms (Likens et al., 1998; Likens & Lambert, 1998). Competition between aluminum and calcium at the roots further decreases levels of available calcium within tissues, while increasing aluminum damages plasma membranes, cell walls, DNA, and increases the burden of oxidative stress. Such nutrient interactions and their broader consequences on physiology and ecology are studied at the Hubbard Brook Experimental Forest (HBEF), a Long Term Ecological Research (LTER) site. It was here that acid deposition was first discovered in North America (Likens & Bormann, 1974) and continues to be studied through the Nutrient Perturbation (NuPert) program (Berger et al., 2001). It provides a replicated high elevation natural ecosystem to examine current, future, and past soil conditions, and has been the site of several key studies on native species, including *A. saccharum, A. balsamea, F. grandifolia*, and *P. rubens*. At HBEF, no studies to-date on *A. saccharum* have focused at the genomic level, where variation in gene expression or signs of adaptation among gene families may be more immediately informative in these slow-growing organisms. For such analysis, a high-quality, chromosomal-length genome is necessary to more accurately detect these forms of variation.

Genomic resources necessary to guide *Acer* conservation are very limited. Only three genomes exist to-date: *A. truncatum* (purpleblow maple), widely distributed across East Asia (Ma et al., 2020), and the endangered *A. yangbiense* and *A. catalpifolium*, native to the Yuhan Province (J. Yang et al., 2019) and West China (Yu et al., 2021), respectively. Here, were present the first two North American *Acer* genomes, *A. saccharum* and *A. negundo*. With these chromosome-scale references, we describe differences in genomic characteristics that may reflect their alternative tolerance strategies. We conducted a differential expression study with stem tissue from *A. saccharum* individuals from HBEF in order to identify genes that may be involved in aluminum response and calcium deficiency. Identification of key processes in the expression study helped to provide focus to the following analysis of comparative gene family dynamics. Together, these approaches highlighted families associated with various abiotic stress responses, including those that also have significant dynamics or novel isoforms in *A. saccharum* or *A. negundo* relative to other broadleaf tree species. And it allowed investigation of the effects of calcium availability at the molecular level, a significant factor associated with maple decline.

## Results

### Genome size estimation and quality control

*A. saccharum* DNA sequencing resulted in 63 Gb of PacBio data with an N50 of 21 Kb (max read length 94 Kb) and Illumina PE data totalling 225 Gb. *A. negundo* was similar with 61 Gb of PacBio data, an N50 of 17 Kb (max read length 95 Kb), and 223 Gb of Illumina PE data. Genome size estimation using short reads resulted in smaller than expected estimates (Contreras & Shearer, 2018; Leitch et al., 2019) at 636 Mb and 319 Mb for *A. saccharum* and *A. negundo*, respectively Figure S1. Using the short-read estimations of genome length, DNA sequence read coverage was high, with long reads at 111x and 141x and short reads at 180x and 208x for *A. saccharum* and *A. negundo*, respectively (Table S1). RNA sequencing of the reference individuals resulted in 61 M reads for *A. saccharum* (92% mapped) and 62 M for *A. negundo* (93% mapped). RNA sequencing of samples for the differential expression study resulted in 207 M reads with mapping rates that ranged from 82 to 92%.

### Genome assembly

Testing of multiple assembly approaches found the FALCON/Purge Haplotigs assembly to be the most contiguous and closest to *A. saccharum’s* estimated size (Figure 2). Statistics for the set of primary contigs from FALCON did not change much between the assembly, unzip, and polishing stages (File S1). The final total length for *A. saccharum* settled at 970 Mb, with an N50 of 691 Kb and 2549 contigs. These last two statistics indicated better contiguity compared to the other assemblers tested. The associated haplotype contigs rose from 109 Mb after assembly, to 320 Mb after unzipping, and dropped to 264 Mb after polishing. Removal of under-collapsed haplotypes reduced the genome size to 668 Mb across 1210 contigs with an N50 of 951 Kb. BUSCO (embryophyte) reported 94.8% complete, but with a somewhat high percentage of duplication (11.7%).

**Figure 2.**
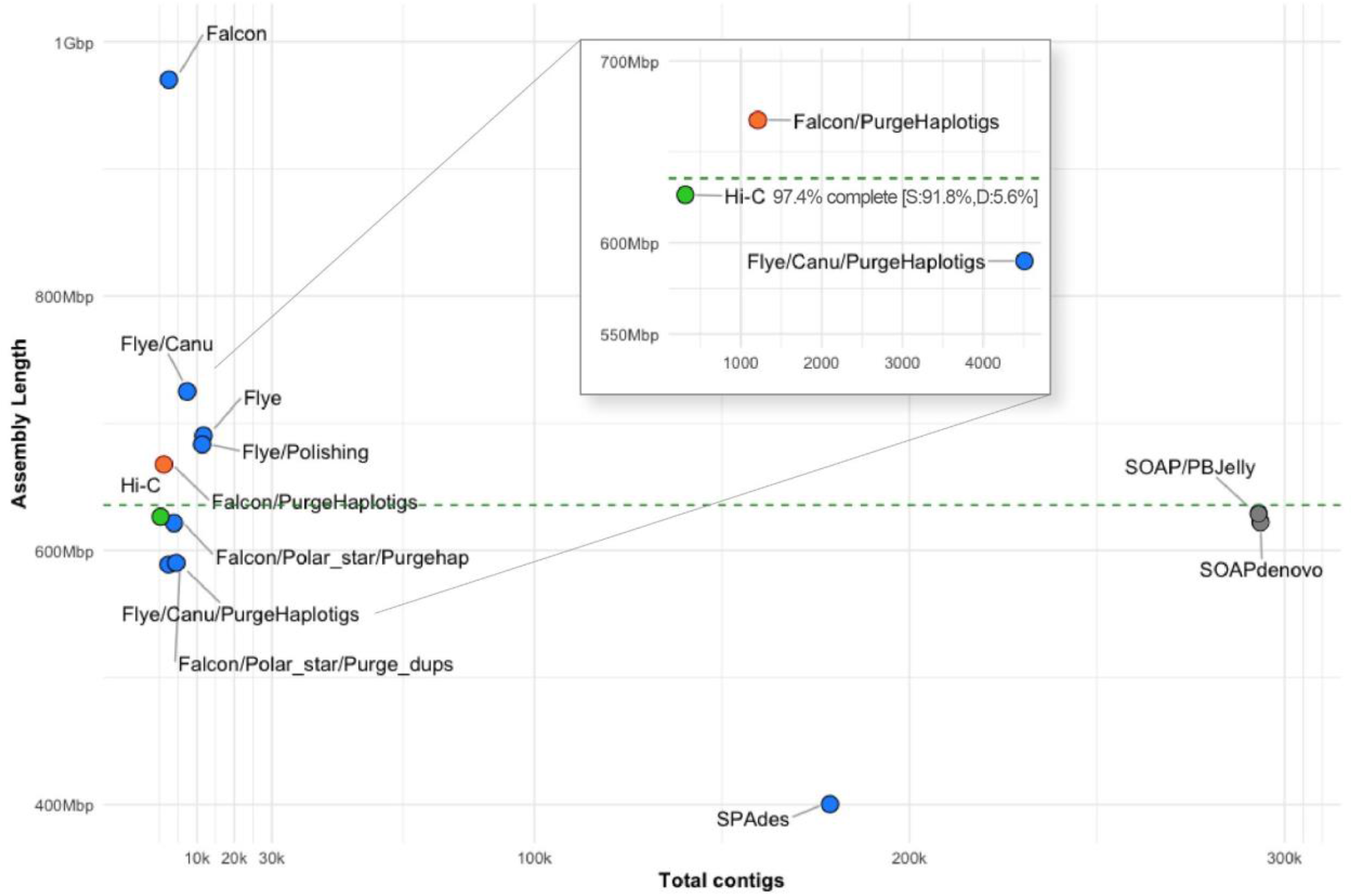
Results of assembly testing with *A. saccharum*, comparing fragmentation in terms of total contigs versus assembly length. The dashed line represents the estimated genome size. Gray dots are short-read assemblers, shown as highly fragmented. Blue dots are long-read tests of assembly workflows. Canu refers to the use of reads error-corrected by the Canu pipeline. The red dot is the selected draft assembly, and the green dot shows scaffolding results following Hi-C. Detailed assembly statistics are available in File S1. https://gitlab.com/PlantGenomicsLab/AcerGenomes/-/blob/master/acer/supplemental/figures/assemblycomparisongraph_busco.jpg

The FALCON primary assembly for *A. negundo* was also consistent in size across pipeline stages, with a total length matching estimates at 481 Mb across 1481 contigs with an N50 of 625 Kb. Removal of haplotype duplication from the primary assembly decreased the overall length to 442 Mb, number of contigs to 1063, and increased the N50 to 700Kb. The BUSCO score was 94.1% with only 5.8% duplication.

### Hi-C scaffolding

The FALCON assembly was selected over Flye due to the substantially more contiguous assembly it produced for *A. saccharum*, though it should be noted that the statistics for both assemblers were comparable for *A. negundo*. Hi-C reads provided 65x coverage of the *A. saccharum* genome. The final assembly was 626.33 Mb in 388 scaffolds with an N50 of 45.72 Mb and GC% of 35.7%. The 13 pseudo-chromosomes represented 97% of the genome length, and BUSCO (embryophyte) scores were 97.7% complete with 3.0% duplicate, 0.7% fragmented, and 1.6% missing.

*A. negundo* Hi-C reads provided 100x coverage and the FALCON assembly was used for scaffolding due to potential mis-assembly in the Flye version (Figure S2). The final assembly was 442.39 Mb in 108 scaffolds with an N50 of 32.30 Mb and GC% of 34.1%. The thirteen pseudo-chromosomes represented 99.74% of the total length. BUSCO embryophyta scores were 97.4% complete with 5.6% duplicate, 0.9% fragmented, and 1.7% missing. (File S1). Earlier work based on low coverage sequencing postulated higher GC content for both *Acer* (Staton et *al*., *2015), and A. negundo* in particular (Contreras & Shearer, 2018), but whole genome sequencing supports their place at the lower range among angiosperm plants (Trávníček et al., 2019).

### Genome annotation

Annotations for *A. saccharum* resulted in 40,074 gene models of which 8,765 were monoexonics verified by the presence of a protein domain and start and stop codons. Transcriptome comparison based on 15,234 transcript loci supported 13,997 gene models. Functional annotation was applied via similarity search or gene family assignment for 35,304 models. *A. negundo* had 30,491 genes, 5,558 of which were monoexonic, and 16,168 transcript loci supported 14,682 of the *de novo* models. Functional annotations were determined for 27,077 of the models. (File S2). *A. saccharum* repeat content was at 64.4% while *A. negundo* was 58.6%.

### Whole genome duplication and Acer synteny

Categorization of putative paralogs revealed a higher percentage of each type in *A. saccharum* relative to *A. negundo*. Plots of Ks distribution for WGD genes in syntenic regions show a single clear peak at a Ks range consistent with the core eudicot WGT reported in other species using the same pipeline (Figure 3a). *A. yangbiense* does not have a recent WGD, and when compared to *A. saccharum* which had an additional small recent peak, further investigation identified small blocks of collinearity, a minimum of five genes in palindromic or tandem arrangements. These blocks are predominantly located on a few scaffolds and are not reflective of the general distribution typical of WGD (File S3). Macrosynteny analysis found that *A. negundo* and *A. yangbiense* are syntenic, while comparisons between each of these and *A. saccharum* revealed a large-scale translocation where two chromosomes from *A. negundo*, including the largest, are split with sections exchanged to form two different chromosomes in *A. saccharum* (Figure 3b, Figure S4).

**Figure 3.**
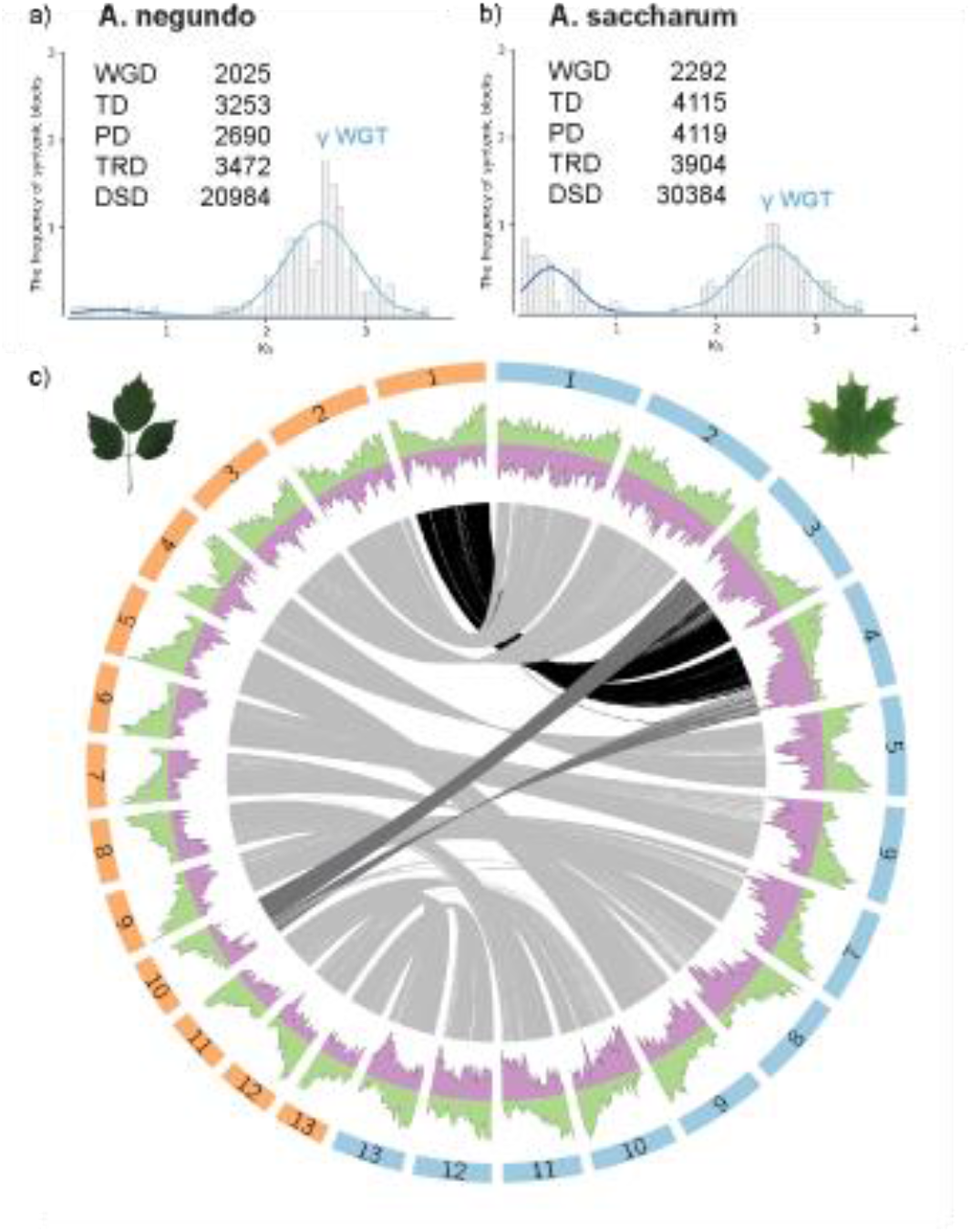
Ks distribution for WGD synteny blocks with a summary of duplication types in (a) *A. negundo* and (b) *A. saccharum*. Abbreviations for categories of duplication: WGD, whole genome duplication; TD, tandem duplication; PD, proximal duplication; TRD, transposed duplication; DSD, dispersed duplication. (c) Circos plot of the thirteen chromosomes ordered largest to smallest for *A. negundo* (orange bars) and *A. saccharum* (blue bars) with distributions of gene density (green) and transposable element frequency (purple). Syntenic regions are linked in gray with darker shades to visually highlight larger recombinations. https://gitlab.com/PlantGenomicsLab/AcerGenomes/-/blob/master/acer/supplemental/figures/acsa_acne_manuscript_circos.png

### Expression analysis of A. saccharum aluminum and calcium treatments

The final annotated genome for *A. saccharum* served as a reference for the expression study. In total, there were 245 unique differentially expressed genes with 181 informatively described by sequence similarity descriptors. Of those with no similarity match, four were completely novel with no identifiable protein domain. Initial analysis produced six up and nine downregulated genes comparing the aluminum to calcium treatments, and the other pairwise comparisons had similarly small totals. Clustering of the expression results showed season had a strong effect (Figure S3), so the analysis was repeated for each season individually to remove this variable from treatment comparisons. For brevity, abbreviations are used according to the following definitions: All, across seasons; Fa, fall; Sp, spring; Al, aluminium; Ca, calcium; and Un, unamended. FaAl to FaCa had 26 upregulated and 17 down, FaAl to FaUn had 7 up and 12 down, and FaUn to FaCa had 28 up and 27 down. SpAl to SpCa had 39 up and 19 down, SpAl to SpUn had the greatest number with 33 up and 41 down, and SpUn to SpCa had 28 up and 11 down (Figure 4, File S4).

**Figure 4.**
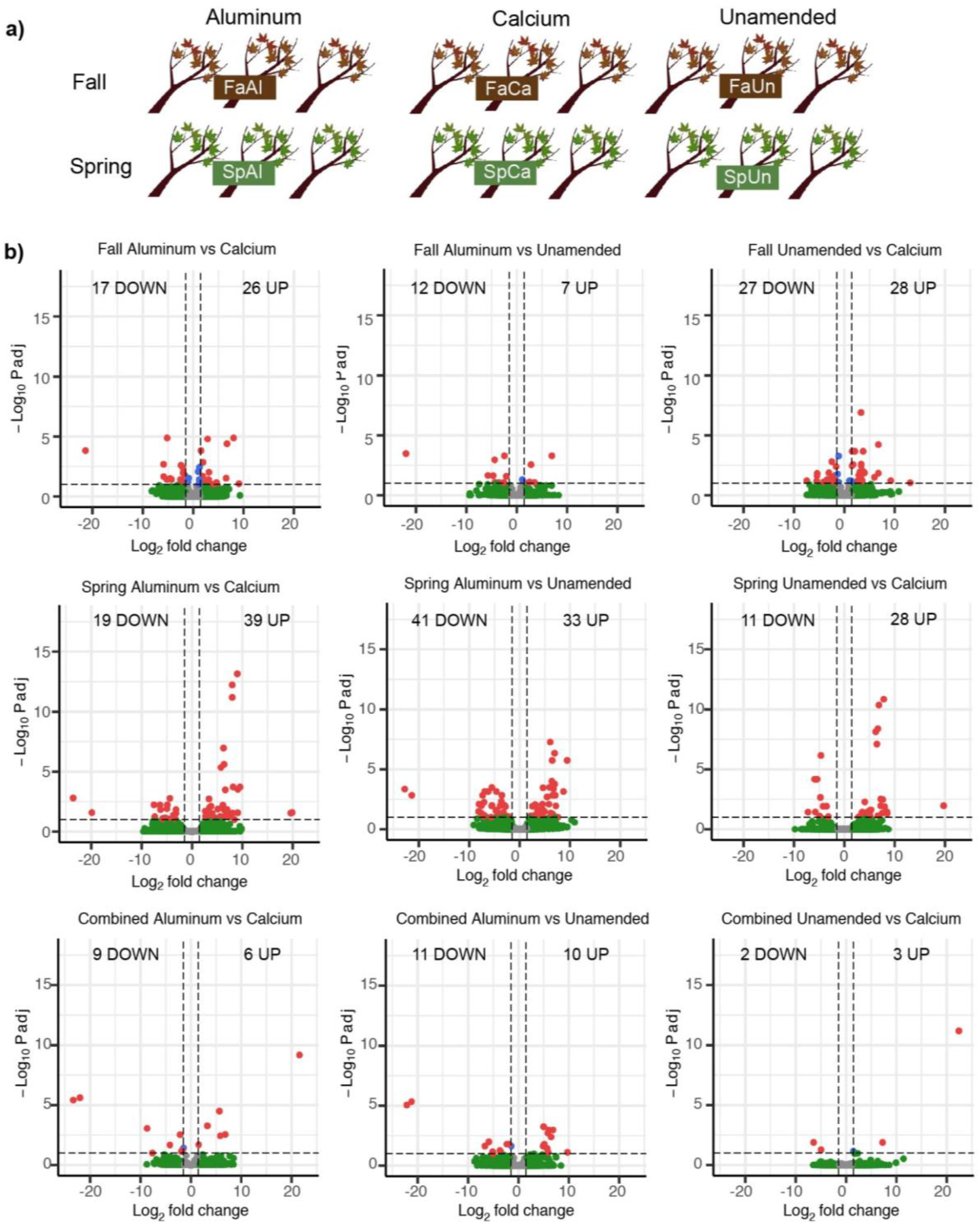
a) Differential expression study design showing number of samples collected in fall and spring from treatment plots at the Hubbard Brook Experimental Forest, Nutrient Perturbation study. b) Differentially expressed genes (up and downregulated) for each treatment and season comparison. Charts display both significance and relative expression denoted as log-fold change. Dotted lines indicate thresholds of significance (0.1 p-adjusted, 1.5 log_2_ fold change). https://gitlab.com/PlantGenomicsLab/AcerGenomes/-/blob/master/acer/supplemental/figures/hbefplots.jpg

There were only two instances where the same gene was found in both separate seasonal analyses. Transcription factor-like E2FE was upregulated in both unamended and aluminum treatments in fall, spring, and all. It had the most isoforms expressed, and was also the most highly upregulated for fall (8-fold). E2FE represses endoreduplication, reducing this type of growth in response to stress (Hendrix et al., 2018). The second instance was disease resistance protein At4g27190-like, expressed primarily in SpAl, but also FaUn and FaCa. Another At4g27190 isoform was upregulated in SpCa and SpAl, but downregulated in SpUn.

GO enrichment results for fall showed sugar and carbohydrate transmembrane transporter activity upregulated in the FaUn compared to FaCa (Table S3). Several sugar transporters, SWEETs and ERD6-like, were seen in both seasons, though more often in fall. There were two SWEET15 (tandem duplicates), one upregulated in FaUn to FaCa, and SWEET15-like upregulated in SpUn relative to both SpCa and SpAl (7.6-fold). SWEET2a was upregulated in FaAl to FaCa. Three different ERD6 were upregulated in FaUn with a fourth in SpAl. Triterpenoid biosynthetic process was also enriched FaUn to FaCa, supported by two downregulated beta-amyrin synthase genes, one of which was significantly downregulated even further in FaAl to FaUn (5.4-fold). Other strongly differentiated genes included accelerated cell death 6-like (ACD6) upregulated in FaCa (5-fold) relative to FaAl. It is associated with flavin-containing monooxygenase 1 (FMO1), the most highly upregulated DEG in FaUn (22-fold) and FaCa, both relative to FaAl.

In spring, DEGs upregulated in SpUn compared to SpAl were enriched in processes related to biotic and abiotic stress, including gene ontology terms for defense response, and acid, oxygen-containing, and antimicrobial response. There were six disease resistance genes, all largely in SpUn and SpCa compared to two in fall. Also present in large numbers, were serine/threonine kinases, including LRR receptor-like, with seven out of twelve of these in the FaUn. Heat shock proteins were also common, though more equally split between the two seasons and shifted toward unamended or aluminum. There were two copies of ITN1, involved in salicylic acid signaling, one of which was the most upregulated DEG in SpCa to SpAl (23.6-fold). In addition to direct stress response, there was an interesting increase of expression in three Holliday junction resolvases, and a lamin-like gene, which was the second highest DEG in these comparisons (19.7-fold). It can play a role in nuclear membrane integrity, chromatin organization, and gene expression (Hu et al., 2019).

#### Metal tolerance via ligation, sequestration, and transport

In SpAl, upregulation of both metallothionein-like 3 (6.6-fold), and aluminum-activated malate transporter (ALMT) 10 (4.5-fold) was observed. An EH domain-containing gene (4.6-fold) is involved in endocytosis, vesicle transport, and signal transduction (Naslavsky & Caplan, 2005), and several cytoskeletal related genes were seen such as kinesin in fall (6.8-fold), and myosin-6 in spring (8-fold).

In addition to the sugar transporters, ABC transporters were present as multiple isoforms in FaUn, and one was the most upregulated DEG in FaUn compared to FaCa (13-fold). Two were downregulated in FaAl compared to FaCa, but still elevated in FaUn, and were from family C which contains pumps for glutathione S-conjugates, which have been shown to remove cadmium in *Arabidopsis* (Tommasini et al., 1998). Additional transport included upregulation of ATPases across seasons, cyclic and mechanosensitive ion channels in spring and fall, respectively, and a K+/H antiporter upregulated in FaAl.

#### Calcium dependent proteins

Three calcium-transporting ATPases were upregulated in either FaCa or FaUn along with an unspecified plasma membrane ATPase. Ca-based external signal relay mechanisms upregulated in SpCa included a cyclic nucleotide-gated ion channel, G-type lectin S-receptor like serine/threonine-protein kinases, and a glutamate receptor (5.5-fold) (X.-L. Sun et al., 2013; Toyota et al., 2018). A calcium-dependent protein kinase (4.5-fold) was upregulated in FaUn. Calmodulin-binding proteins (60d at 5-fold) were up-regulated in both SpAl and SpCa, and are associated with salicylic acid synthesis and immunity in *Arabidopsis* (L.-S. Li et al., 2021).

#### Reactive oxygen species response

Antioxidant and redox genes included glutathione peroxidase, upregulated in FaUn, and glutathione S-transferase, upregulated in both FaCa and SpAl. Two cytosolic sulfotransferases and six thioredoxin genes were upregulated in SpAl. Other redox related genes include a highly expressed epoxide hydrolase (9-fold) and a carotenoid cleavage dioxygenase (4.6-fold) in FaAl; and peroxidase, aldehyde dehydrogenase (6.7-fold), and germin in SpAl. Ascorbate-dependant oxidoreductase SRG1 was upregulated in SpUn, cytochrome P450 71D11 (a monooxygenase) in SpAl, and zinc finger (C2H2 type) in SpUn.

#### Cell wall and membrane integrity

MDIS1-interacting receptor kinase, upregulated in FaUn relative to FaAl, and expansin, FaUn to FaCa, are associated with the cell wall. Two ADP-ribosylation factor GTPase-activating, one of which is the most highly upregulated gene in spring (19.9 fold-change SpAl to SpCa), assists with cell signaling by recruiting cargo-sorting coat proteins at the membrane and regulating lipid composition in support of development and defense (Donaldson & Jackson, 2011).

#### Hormone crosstalk

Auxin was indicated by three WAT genes (seen in each season, greatest at 7-fold in SpAl to SpCa) and indole-3-acetic acid (highest of AllAl to AllUn at 9.7-fold). Two ethylene synthesis genes were present: 1-aminocyclopropane-1-carboxylate oxidase, FaAl to FaCa, and methylthioribose kinase, SpAl to SpCa. Ent-kaurenoic acid oxidase is related to brassinosteroid homeostasis and gibberellin biosynthesis (Helliwell et al., 2001) and was present in SpAl to SpUn (7-fold), and obtusifoliol 14*α*-demethylase, which mediates brassinosteroid synthesis, was upregulated SpUn to SpCa (Xia et al., 2015). Jasmonic acid activity was upregulated in FaCa, negative regulation of cytokinin was up in AllAl to AllUn, and ABA-induced HVA22 which inhibits gibberellin and is possibly involved in vesicular traffic, was in SpAl to SpCa (8-fold) as well as AllAl to AllCa.

### Gene family evolution with expression study integration

By leveraging 22 high-quality plant genomes, gene family dynamics between *A. negundo* and *A. saccharum* revealed distinct characteristics. Comparisons among the plant proteomes resulted in 20,234 orthogroups with a mean size of 26.4 genes. Of these, 4,262 were shared by all species, and 79 were single-copy. 88.5% of 603,640 genes were contained in orthogroups, and 0.7% were species-specific. All species had at least 80% of their genes contained in orthogroups with the exception of *Ginkgo biloba, Nymphaea colorata*, and *Oryza sativa* (Table S4).

All three *Acer* shared 11,156 orthogroups. *A. negundo* and *A. saccharum* had the largest overlap, *A. saccharum* and *A. yangbiense* had the second largest overlap, and *A. saccharum* had the most unshared groups. (Figure 5b). Comparing *Acer* against the other woody angiosperms (*B. pendula, C. papaya, C. clementina, C. sinensis, E. grandis, J. hindsii, J. regia, P. vera, P. trichocarpa, P. persica, Q. lobata, Q. robur, T. grandis*), 728 *Acer* orthogroups were expanded, 14 were contracted, 1992 were novel, and *Acer* was estimated to be missing from 202 orthogroups. To clarify, these may not be fully absent, but didn’t have representation in orthogroups including those that were more lineage specific. Comparing the Sapindales to the other trees resulted in 340 expanded, 4 contracted, 2788 novel, and 160 missing. Dynamics in common between the two *Acer* included 270 expanded groups, 1 contracted, 0 novel, and 247 missing (File S5).

**Figure 5.**
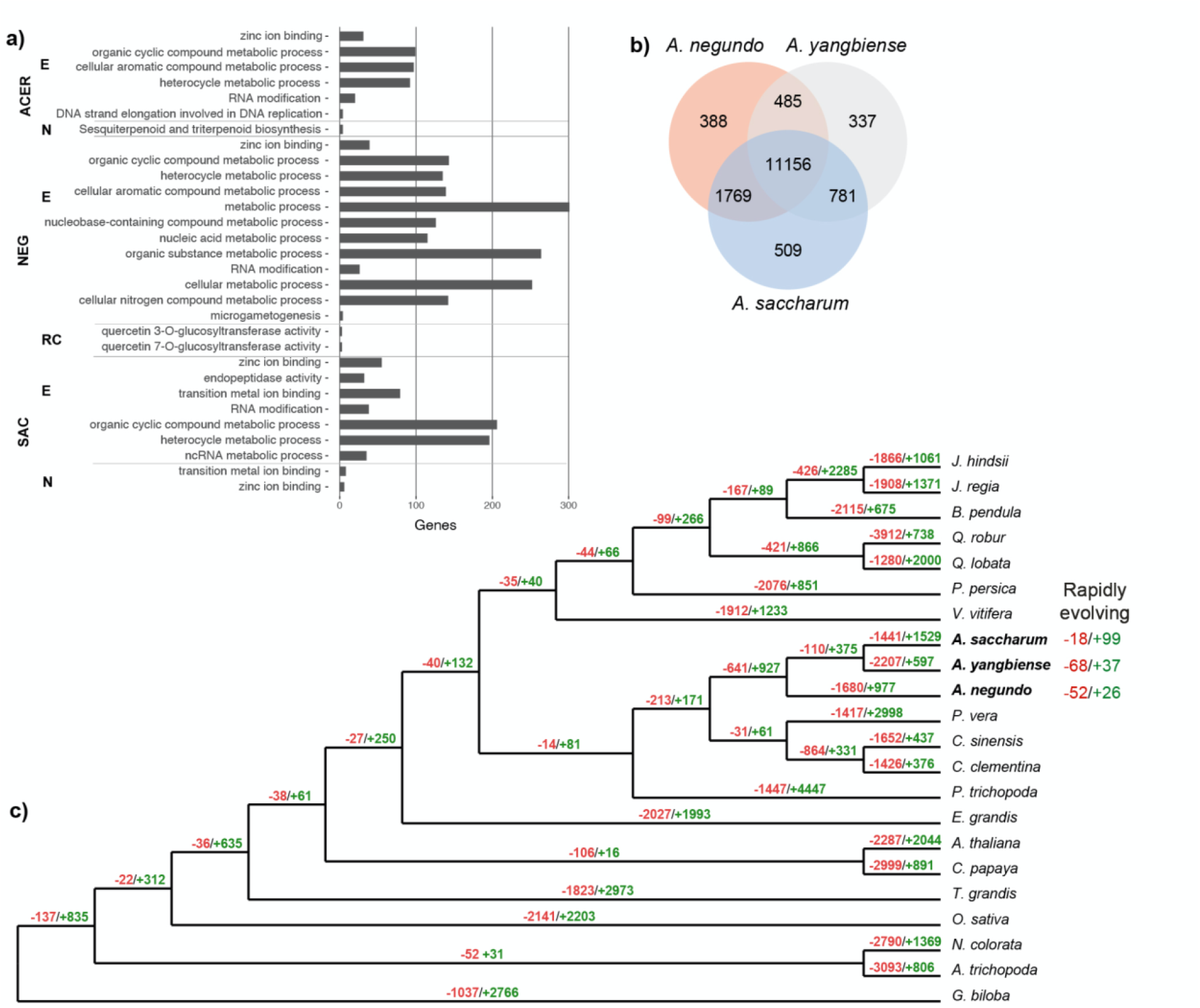
a) Gene ontology enrichments for *Acer* (all three species combined), *A. negundo*, and *A. saccharum*. Abbreviations for gene family dynamics: E, expanded; N, novel; RC, rapidly contracting. b) Total gene families, shared and unique, among the *Acer*. c) Reconstructed gene tree showing contracted gene families in red and expanded in green. https://gitlab.com/PlantGenomicsLab/AcerGenomes/-/blob/master/acer/supplemental/figures/cafetreevennGO.jpg

*Acer* families expanded among the woody angiosperms were enriched for RNA modification, DNA replication strand elongation, and processes of organic cyclic compound, cellular aromatic compound, and heterocycle metabolisms. Novel genes were highly enriched for cell periphery localization and marginally for sesquiterpenoid and triterpenoid biosynthesis (Figure 5a, Table S5). When focusing on absent gene families, none were found to be missing exclusively in Sapindales, which includes the two *Citrus* species and *Pistacia vera*. A total of two families were absent in all *Acer*, phosphatidylcholine transfer protein-like and cellulose synthase interactive 3, but were present in all other species.

Several interesting gene families more novel to *Acer* overlap with the HBEF DEGs. There are twenty orthogroups associated with disease resistance At4g27190, two seen as DEGs, and the specific families containing these DEGs are larger for *A. saccharum* with one novel to the *Acer*. Two additional non-DEG families are rapidly expanding in *A. saccharum*, with one of these also expanding in *A. negundo*, but both contracting in *A. yangbiense*. In fall, another more novel DEG ACD6-like belongs to a family with limited species membership (Sapindales and *V. vinifera*). Compared to *Arabidopsis* ACD6, which has two 3-repeating ankyrin domains, the *A. saccharum* ACD6-like had varying ankyrin positions, as do other members of this family. A third example, seen only in spring, acetyl-coenzyme A synthetase (ACS) is a member of a novel *Acer* family consisting of 1 *A. negundo*, 6 *A. saccharum*, and 18 *A. yangbiense*. Comparison with *A. negundo* and ACS isoforms reveals the *A. saccharum* gene contains a longer ACL domain, with only ∼67% query coverage and ∼43% percent identity to *A. negundo* which is much more similar to other ACS (87-90% length; ∼88% identity).

Within the broader set of species, *A. saccharum* gene families were characterized by more expansion, with 1827 expanded, 18 contracted, 127 novel, and 511 absent, and more rapidly expanding, with 99 compared to 18 contracting. *A. negundo* had 1068 expanded, 23 contracted, 89 novel, and 558 absent orthogroups. Rapidly contracting families were greater in this species with 52 compared to 26 rapidly expanding (Figure 5, File S5, File S6).

#### A. saccharum gene families

Functional enrichment of *A. saccharum* in the full species comparison revealed that expansions were processes of ncRNA metabolism, RNA modification, organic cyclic compound metabolism, heterocycle metabolism, and intracellular membrane-bound organelle localization (Table S5). Almost half of the *A. saccharum* families had limited annotation information, due to either missing descriptors or uncharacterized protein matches. Relative to *Acer, A. saccharum’s* expanded families are enriched in a larger list of stress response associated functions that are fairly specific, including water deprivation, hypoxia, salinity, heat, cold, xenobiotic, nematode, karrikin, acid chemical, and hormone (File S13). Other significant processes include regulation of indolebutyric acid stimulus (auxin family), RNA splicing, chloroplast RNA processing, phospholipid translocation, brassinosteroid homeostasis, lignin synthesis, microsporogenesis, phenylpropanoid biosynthesis, cadmium ion transmembrane transport, cyclin-dependent serine/threonine kinase, and calcium-transporting ATPase. Rapidly expanding families are associated with various biotic and abiotic responses, such as fungal, salt stress, and xenobiotic response (File S6). Interesting genes include patatin-like 2, involved in membrane repair via removal of lipids modified by oxidation (W.-Y. Yang et al., 2012) and ALP1 negative regulation of polychrome group chromatin silencing (Liang et al., 2015). Those that overlap with HBEF DEGs include disease resistance At4g27190 and DSC1, FMO1, rapidly expanding SRG1, and rapidly contracting disease resistance At1g50180.

Compared to other *Acer*, contracted families are enriched for pollen wall assembly, extracellular matrix assembly and organization, chlorophyll binding, NADH dehydrogenase (ubiquinone) activity, DNA-directed 5’-3’ RNA polymerase activity, programmed cell death, and myb-like transcription factors (File S11, File S12). Genes absent in *A. saccharum* but present in all other species total 28 (File S15), including red chlorophyll catabolite reductase (ACD2) and S-adenosyl-L-homocysteine hydrolase (HOG1), which is necessary to hydrolyze the by-product of the activity of S-adenosyl-L-methionine-dependent methyltransferase, and one of these was also absent. The apparent absence of HOG1 requires further investigation as mutants display a number of problematic phenotypes and variants display association with fiber length in *P. tomentosa* (Du et al., 2014).

#### A. negundo gene families

In the broad comparison of species, *A. negundo* expanded families were enriched in RNA modification, microgametogenesis, and metabolic processes of nucleobase-containing compound, organic cyclic compound, heterocycle, and cellular aromatic compound (Table S5). The small number of rapidly expanding families were by far mostly uncharacterized proteins or missing sequence similarity descriptions with only four out of 26 genes having a description, including glutathione-S-transferase, two disease-resistance proteins, At4g27190-like and At5g66900, a receptor-like 12, and additional functional descriptors such as E3 ubiquitin-protein ligase, LRR receptor-like ser/thr kinase, and more (File S6). Relative to other *Acer, A. negundo’s* expanded families are enriched in a short list of specific stress response including UV, UV-B, radiation, bacterium, cadmium, metal ion, drug, chemical, and osmotic stress (File S9, File S10). Other processes include proanthocyanidin biosynthesis, lignin synthesis via cinnamyl-alcohol and sinapyl-alcohol dehydrogenase, starch metabolism and glucan catabolism, error-prone translesion synthesis, and other DNA damage response and repair. Processes related to reproduction were present, especially pollen development. For example, decreased size exclusion limit 1 (DSE1, aka aluminum tolerant 2 (ALT2)) is a transcription factor that regulates the size of molecules that can travel through plasmodesmata, as channel aperture is not static, changing in response to stress and decreasing during embryo development (Xu et al., 2012). DSE1 is single copy in all species with expansions only in *A. negundo, Q. rober*, and *T. grandis*.

Contractions relative to other *Acer* include transcription by RNA polymerase III, chloroplast RNA processing, and lignin biosynthetic processes (File S7, File S8). Rapidly contracted families were enriched in quercetin 3-O- and 7-O-glucosyltransferase activity (Table S6). Examples of contracted families include 7-ethoxycoumarin O-deethylase, which metabolizes a wide range of xenobiotics (Robineau et al., 1998), and disease resistance-like protein DSC1, both rapidly expanding families in *A. saccharum* (File S6). There are 23 genes absent in *A. negundo* that are present in all other species (File S14). Several of these are curious, as they appear to be required components of important processes, such as AUGMIN subunits 2 and 7 that help form a complex that plays a role in spindle microtubule generation (Tian & Kong, 2019), and cell division cycle 45-like, which is required for meiosis in *Arabidopsis* (Stevens et al., 2004). The absence of formamidopyrimidine-DNA glycosylase is also interesting as it is involved in base excision repair of DNA damage, a notable area of specific enrichments described below.

## Discussion

*Acer saccharum* and *A. negundo* differ in species distribution and tolerance to abiotic stressors, and their genomes reveal support for their contrasting life histories. *A. saccharum* has slow growth until release of canopy coverage, doesn’t achieve reproductive maturity until 40 years of age, and is able to live 400 years. It requires high nutrient soils, prefers mesic environments, and is tolerant of cold, but not salinity. Its range crosses a more narrow latitudinal gradient, but a stronger elevational one, where *A. negundo* tends to be more limited. While both genomes are small in size and diploid, *A. saccharum* is 42% larger, contains twice as many transposable elements, and 38% more gene duplications, many of which are very recent. Its gene families tend to be larger, more diverged, and undergoing rapid expansion compared to *A. yangbiense* or *A. negundo*.

In contrast, *A. negundo* grows quickly, is able to reproduce after only five years, and has a relatively short lifespan of 60 years. It is tolerant of a range of conditions and is widespread throughout North America. *A. negundo* is considered an aggressive invasive in Europe, South Africa, parts of Asia and North America, and it rapidly dominates disturbed habitats (CABI, 2021). It is sexually dimorphic and both native and invasive conspecifics are highly plastic in growth, leaf phenology, and physiological traits such as maximum assimilation rate, nitrogen content and photosynthetic efficiency (Dawson & Ehleringer, 1993; Lamarque et al., 2015). Relative to native heterospecifics, invasive populations of *A. negundo* maximized growth in high light and nutrient conditions, with dramatically reduced growth and survivorship in deficient environments (Porté et al., 2011).

The *A. negundo* reference genome is a small diploid with high heterozygosity from the native species range. Macrosynteny between *A. yangbiense* and *A. negundo* is largely collinear and without rearrangement as shown in the two *A. saccharum* chromosomes (Figure S4). Invasive plant species are often associated with smaller genomes and *A. negundo* is characterized by contracting gene families. Traits such as fast growth rate, germination time, stomatal responsiveness, and dispersal ability are cell size or division rate dependent (Pyšek et al., 2018). The greater surface area to volume ratio of small cells, derived from small genomes, reduces the metabolic and signaling requirements, but does not preclude additional growth or activation, thus extending the range of expression, or plasticity, for a wider set of traits (Roddy et al., 2019; Suda et al., 2015). The adaptive potential conferred by polyploidism can also be leveraged for success of invasives, but polyploidism has not been documented in the native or introduced range.

If larger genome size presupposes increased metabolic, transport, and nutrient demands (Pellicer et al., 2018) perhaps *A. saccharum’s* susceptibility to nutrient deficiencies, may be in part due to the extra burden of a larger genome and its particular enrichments. The decrease in soil nutrient availability in its native range over the past decades is at odds with resources necessary to tolerate stressors brought on by a changing environment, and previous adaptations are now maladaptive. From the foundational differences between *A. negundo* and *A. saccharum* in morphological, physiological, and genomic characteristics, we turn to the integration of gene family dynamics and expression data from a study on nutrient stress to further elucidate the contrasting strategies of tolerance seen in these species.

### Integration of Acer saccharum expression studies

Integration of gene expression sampling with gene family dynamics highlighted the genetic factors that may influence maple decline and adaptation. HBEF NuPert *A. saccharum* saplings grown in native ecological and environmental conditions within replicated long-term soil treatments provided an opportunity to examine differences in gene expression related to soil acidity, calcium deficiency, and increased aluminum availability (Table S7). Calcium amendment of NuPert plots was designed to represent levels from 50 years ago, before extensive acid deposition in the HBEF range led to the deficient conditions seen in unamended control plots (Berger et al., 2001). Aluminum amended plots further decrease calcium availability via competition for uptake between the two cations. They represent more extreme calcium depletion and an increase in aluminum availability that is observed with soil acidity. Examination of trees on the landscape in these conditions allowed us to study the effects of long-term nutrient stress in conjunction with other life-history processes. Seasonal differences between the differentially expressed genes were striking, revealing the effect of calcium levels on processes occurring at the time of sampling.

### Signaling and metabolic processes are sensitive to lower calcium levels

Calcium is the common limiting element underlying maple decline, and trees improve long-term with the addition of calcium (Moore & Ouimet, 2021). Calcium transport and binding is well studied for its ability to activate and regulate primary components of stress response systems, including internalization of external signals, modulation of cytosolic levels for transient signaling, and initiation and perpetuation of ROS responses, hormone modulation, and pathways of secondary metabolism (Estravis-Barcala et al., 2020). In physiological studies conducted at HBEF, calcium amendments resulted in a 50% increase of calcium in foliar tissues of *A. saccharum*, while non-significantly decreasing aluminum concentrations. Trees in calcium plots devoted more carbon to growth than storage, were better able to flush after a late spring frost, produced more flowers, and increased seed germination (Halman et al., 2013).

Calcium amendment facilitated upregulated gene expression relative to unamended and aluminum treatments in areas related to calcium signaling and senescence activities. In fall, a gene involved in leaf senescence was highly expressed in calcium treatments, and the relative paucity of other differential expression in this season made it all the more interesting. ACD6 is important to both natural and stress response senescence, shown to regulate rosette biomass, chlorophyll degradation, and leaf nitrogen and carbon levels in *A. thaliana* (Jasinski et al., 2021). It is also a factor in nitrogen translocation from leaves to seeds before cell death. ACD6 stimulates calcium signaling in *A. thaliana* and could potentially be an ion channel itself (Zhu et al., 2021), and several calcium dependent signaling and calmodulin binding genes were upregulated in the fall calcium and unamended plots, including the critical calcium-dependent protein kinase 1. The ACD6 isoform expressed here is limited to Sapindales and quite diverged from the *A. thaliana* form. Interestingly, a second senescence gene that is expressed in *Acer rubrum* (Z. Chen et al., 2019) is absent from the *A. saccharum* genome. Red chlorophyll catabolite reductase (RCCR aka ACD2) is a non-green chlorophyll degradation, and while the exact role of RCCR is unclear (Jockusch & Kräutler, 2020), it appears possible that like many senescence related genes, it provides additional fitness advantages (Mach et al., 2001). Timing, duration, and intensity of gene expression are all important during senescence activities leading up to cell death, including adjustment of signaling, growth, metabolism, and nutrient remobilization. The susceptibility of ACD6 to calcium deficiency could potentially create complications in fall that could affect spring growth, reproduction, and ability to respond to stress.

Throughout *A. saccharum’s* native range, when calcium is deficient, additional nutrient imbalances, such as magnesium, phosphorus, potassium, and nitrogen deficiency, as well as high aluminum and manganese concentrations, climatic effects, and biotic stressors such as defoliation, create compounding stress (Bal et al., 2015; Long et al., 2019). Samples from unamended plots show an increase in stress response perhaps reflective of calcium deficiency combined with external stressors. Signs include enrichments of sugar and other transporters in fall samples (Julius et al., 2017; Miao et al., 2017). Spring samples reported upregulation of several putative disease resistance genes, heat shock proteins, leucine-rich repeats proteins and kinases, and signs of oxidative stress.

### Oxidative stress response is high in aluminum amendment conditions

Reactive oxygen species (ROS) are generated as a result of both stress and normal cellular processes, and reactive oxygen can take different forms, with varying toxicities. ROS is reduced by a variety of potential “scavengers” to prevent accumulation of levels that are damaging to lipids, proteins, DNA, and RNA, leading to cell death. ROS production and redox activity vary according to the primary processes of each cellular compartment (i.e., metabolism in mitochondria), and also in response to different stressors or stress combinations (i.e., drought and heat), so it is thought that cells develop complex ROS signatures or hotspots that influence signal transduction and metabolic regulation in nuanced ways (Castro et al., 2021; Choudhury et al., 2017). Aluminum increases peroxidation of lipids in membranes and produces H_2_O_2_ which can participate in retrograde signaling, regulating expression of additional genes (Castro et al., 2021). Aluminum derived oxidative stress causes inhibition of cell growth and is an early signal of aluminum toxicity (Yamamoto et al., 2003).

Antioxidants and redox genes are commonly observed as a response to ROS stress and a combination of these were observed in different seasons and treatments, though more were found in spring, as fits with the trend of increased overall activity in that season. Antioxidants, including thioredoxins, peroxidases, and a germin-like, were all upregulated in the aluminum treatment, implying significant stress response activation. Glutathione system members, which can also act as chelators, are more broadly seen in both unamended and aluminum. A number of antioxidant DEGs were also from expanded families, in particular the significantly expanded thioredoxins H3 and YLS8 and SRG1 (Figure 6). The antioxidant DEGs from this study can be compared with a number of other transcriptomic studies on metal stress in trees where glutathione S-transferase, peroxidases, and thioredoxins were in common. For example, cadmium accumulators *Salix integra* and *Populus x canadensis* ‘Neva’ both expressed superoxide dismutase with glutathione pathway genes and peroxidases, respectively (X. Li et al., 2021; Shi et al., 2016). Nickel stress in resistant versus susceptible genotypes of *Betula papyrifera* found glutathione S-transferase and thioredoxins upregulated in resistant trees (Theriault et al., 2016). *A. rubrum*, a nickel avoider, had root-level expression of superoxide dismutase, but downregulated glutathione S-transferase (Nkongolo et al., 2018) similar to one of the *A. saccharum* DEGs. In comparisons of aluminum treated *Citrus* root tissue expression, peroxidases and germin-like proteins were upregulated. It is interesting that superoxide dismutase was not significantly expressed in *A. saccharum* given that it is from an expanded gene family and seen in several other species. In physiological studies conducted at HBEF, Halman et al. (2013) found *A. saccharum* were under increasing oxidative stress in unamended and aluminum plots. Aluminum treated trees had elevated levels of glutathione reductase activity as measured by spectrophotometric assay. Ascorbate peroxidase activity was also elevated, but this particular peroxidase was not observed in our gene expression study.

**Figure 6.**
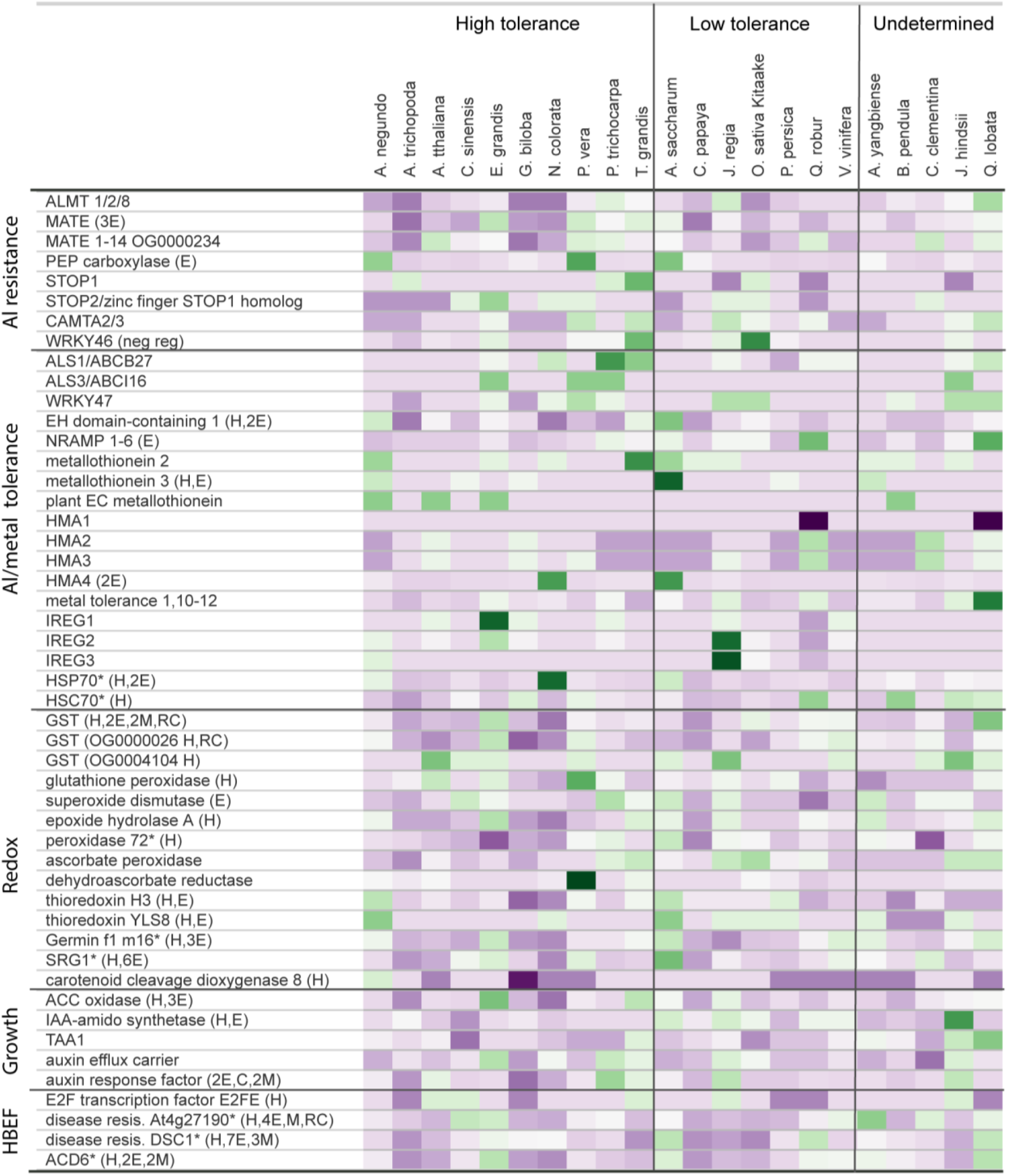
Orthogroup sizes for aluminum tolerance gene families are presented by species. Families were selected for inclusion based on documented aluminum tolerance and/or presence in the HBEF RNA-Seq differential expression results. Color represents the proportion of gene membership per species, with darker purple equating to more contracted families relative to the median, and dark green indicating expansion. (H) Family contains HBEF differentially expressed gene; (E) Expanded in *A. saccharum*; (C) Contracting; (M) Missing; (N) Novel; (*) Rapidly expanding; Categorization of tolerance is according to literature describing aluminum stress phenotypes. The undetermined category contains species where tolerance to aluminum or acidic soils has not been reported. ^1^*B. pendula* is undetermined due to high variability in tolerance by genotype. https://gitlab.com/PlantGenomicsLab/AcerGenomes/-/blob/master/acer/supplemental/figures/genefamily_al_comparisons_manuscript_customnorm.html.jpg

### Aluminum and metal response is minimal in aluminum amendment conditions

Deciphering aluminum response mechanisms can be complicated as aluminum affects several targets within cells, and resistance and tolerance mechanisms can vary widely, even within species. Accumulation of aluminum occurs primarily in the root, binding to pectin in the cell wall, increasing its rigidity and decreasing permeability. Throughout the plant, aluminum can disrupt plasma membrane potential and membrane-bound solute transporters (Kar et al., 2021). Damage to proteins and DNA reduces enzymatic activity, and oxidative stress and hormone signaling from DNA damage can lead to growth inhibition (Sade et al., 2016).

Studies from locations such as Vermont, the Adirondack Mountains, and the Allegheny Plateau, have linked foliar aluminum and calcium levels (Long et al., 2019; Oswald et al., 2018; Sullivan et al., 2013) and stem aluminum levels to branch dieback as seen in maple decline (Mohamed et al., 1997). At HBEF, previous studies examined physiological effects of aluminum with a multi-tissue study on mature *A. saccharum*. Using inductively coupled plasma atomic emission spectroscopy, Halman et al. (2015) found foliar levels of aluminum did not vary much by treatment and all were well below thresholds of toxicity. Calcium treatments caused concentrations of aluminum to drop slightly, while aluminum treatments caused calcium to drop more strongly in dominant than non-dominant trees. The main effects noted were moderate root damage and foliar antioxidant activity (Halman et al., 2013). Within our stem tissue expression analysis, signs of metal response were not clear, but we can detect levels of general stress response expression between the unamended and aluminum treatments.

Genes currently known to be associated with aluminum resistance or tolerance were not expressed in the stem tissue samples, but upregulated genes related to metal transport and sequestration are potentially involved in aluminum remediation (Hasan et al., 2017; Hasanuzzaman et al., 2017). Metallothionein 3 was highly expressed in the combined season aluminum comparisons, though unevenly distributed throughout the replicates. Similar to other metallothioneins, it works by ROS scavenging and metal ion homeostasis via chelation. (Hasan et al., 2017). In salt-tolerant *Oryza sativa*, this gene responds to cadmium, salinity, and oxidative stress (Mekawy et al., 2018).

Most of *A. saccharum’s* aluminum resistance gene families are modestly sized relative to other species examined, with the exception of PEP carboxylase and a large family of mixed MATEs, with members likely similar to homologs studied in *P. trichocarpa* and *Arabidopsis* (N. Li et al., 2017). Overexpression of either of these increases efflux of organic acids at the root (Begum et al., 2009). *A*.*saccharum* had slightly more expansion in resistance families than the other *Acer*, while *A. negundo* had the least. Given the limited number of DEGs indicating metal toxicity, and the non-toxic aluminum tissue concentrations observed in physiological studies, it seems likely resistance mechanisms exist at the root in *A. saccharum*. Comparing aluminum/metal tolerance gene families, *A. yangbiense* had the fewest expanded and there are no reports on aluminum tolerance for this species. *A. negundo* and *A. saccharum* were not notably different with the exception of the metallothionein 3 family, which was rapidly expanding, much larger in *A. saccharum* relative to other broadleaf tree species, and observed in the expression results for both seasons. Expanded resistance and tolerance families were seen more frequently in high-tolerance species such as *P. vera, P. trichocarpa, T. grandis, Q. lobata*, and *E. grandis*, though the mix of families varies (Figure 6; Sork et al., 2016; Tuskan et al., 2006; Zeng et al., 2019; Zhao et al., 2019). Gene family size is only a potential factor in response and it does not necessarily serve as a proxy for expression. The redox families selected for comparison are primarily limited to those containing DEGs, and in this case there is good overlap between gene family size and expressed redox genes. Many were larger in *A. saccharum*, but *A. negundo* was also enriched for a different set of redox families.

### Indications of growth reduction and DNA damage in response to aluminum

Strong upregulation of transcription factor E2FE was notable as it is one of only two genes seen in both seasons. E2FE represses endoreduplication (Liu et al., 2019), and while endoreduplication can be induced by certain types of stress such as drought and salinity (Lang & Schnittger, 2020), cadmium toxicity was observed to reduce endoreduplication and cell division in *A. thaliana* leaves (Hendrix et al., 2018). With cadmium, pathways that delay cell cycle progression due to oxidative DNA damage were upregulated along with E2FE/DEL1. Aluminum is known to cause double-stranded breaks which requires homology-dependent recombination repair (P. Chen et al., 2019). In *A. saccharum*, DNA damage might be indicated by upregulation of several Holliday junction resolvases which help repair double stranded breaks using homologous recombination (Bauknecht & Kobbe, 2014). Patterns of expression point toward oxidative DNA damage from aluminum or another source, inducing repression of endoreduplication similar to cadmium.

The hormone Indole-3-acetic acid (IAA)-amido synthetase was the most highly upregulated gene in the combined season comparison of aluminum treatment to unamended. The gene family for this DEG is novel to *Acer* and significantly expanded in *A. saccharum*. IAA is the most abundant auxin in plants, and IAA-amido synthetase helps with its homeostasis by leading to IAA-amino acid conjugates (Ludwig-Müller, 2011). Increased expression of IAA-amido synthetase produces highly reduced growth forms, altered architecture, and shortened roots (C. Ding et al., 2021; M. Sun et al., 2020). In *O. sativa*, over-expression of IAA-amido synthetase resulted in improved drought tolerance (Zhang et al., 2009), fungal pathogen response (X. Ding et al., 2008; Domingo et al., 2009) and altered growth. Park et al (2007) observed reduced growth in *A. thaliana* from overexpression of an IAA-conjugating gene (GH3) that was highly linked to biotic and abiotic stress response. They proposed that modulation of auxin is an adaptive strategy that trades reduced growth for enhanced stress resistance.

### Stress response in Acer saccharum and calcium deficiency versus aluminum toxicity

Replicated samples of *A. saccharum* reveal complex expression changes indicating multiple forms of stress response in aluminum treatments and to a lesser degree in unamended plots. Differentially expressed genes, often related to signaling, redox, hormones, and growth reduction, are seen in aluminum or unamended treatments, but not as extensively in calcium, where there is enhancement of other processes, such as senescence and calcium signaling. Ecosystem and environmental conditions at the time of sampling likely had interacting effects and contributed to complexity of expression, but trends by season and treatment were still evident. With this data, we provide preliminary molecular support to the many studies correlating maple decline with calcium-poor, acidic soils. Given the minimal expression of metal tolerance genes and lower level of foliar aluminum seen in dominant sugar maple at HBEF, along with large amount of variability in aluminum response sometimes seen between conspecific genotypes in other species, further transcriptomic studies of root tissue would be beneficial for a better understanding of aluminum resistance in *A. saccharum*.

### Herbivory, DNA damage, and light harvesting enrichment in Acer negundo

Gene family expansion in *A. negundo* was specifically limited to UV, bacteria, metal, cadmium, chemical, and osmotic response rather than the more extensive set seen in *A. saccharum*. A high proportion of proanthocyanidin synthesis related gene families were expanded, which are precursors to condensed tannins that protect against herbivory, bacteria and fungal pathogens, and encroaching of neighboring plants (He et al., 2008). All four families were relatively novel to *A. negundo* with absence in all but one or two other species. Proanthocyanidins also have antioxidant and radical scavenging functions, and have been compared with lignins in terms of pathogen defense mechanisms (Stafford, 1988).

Both *Acer* are enriched in gene families with DNA damage recognition and repair functionality, but *A. negundo* has a greater number, and they are largely categorized as either translesion synthesis or meiosis-related repair. Translesion synthesis is often activated by UV damage. It is an error-prone repair mechanism designed to quickly eliminate lesions at the replication fork that might lead to stalling and double-stranded breaks (Sakamoto, 2019). *A. negundo* has three families in this category expanded relative to *Acer*, two of which are significantly expanded among the full comparison of species. (Nisa et al., 2019). Among *A. negundo’s* DNA damage related enrichment are secondary enrichments that include regulation of leaf, seed, flower development, and post-embryonic structures, response to abiotic and biotic stress, and growth such as cell division and endoreduplication (Figure 7). Too much DNA damage can inhibit growth and development or initiate programmed cell death. Perhaps a more robust response to DNA damage is a factor in *A. negundo’s* plasticity of growth and tolerance.

**Figure 7.**
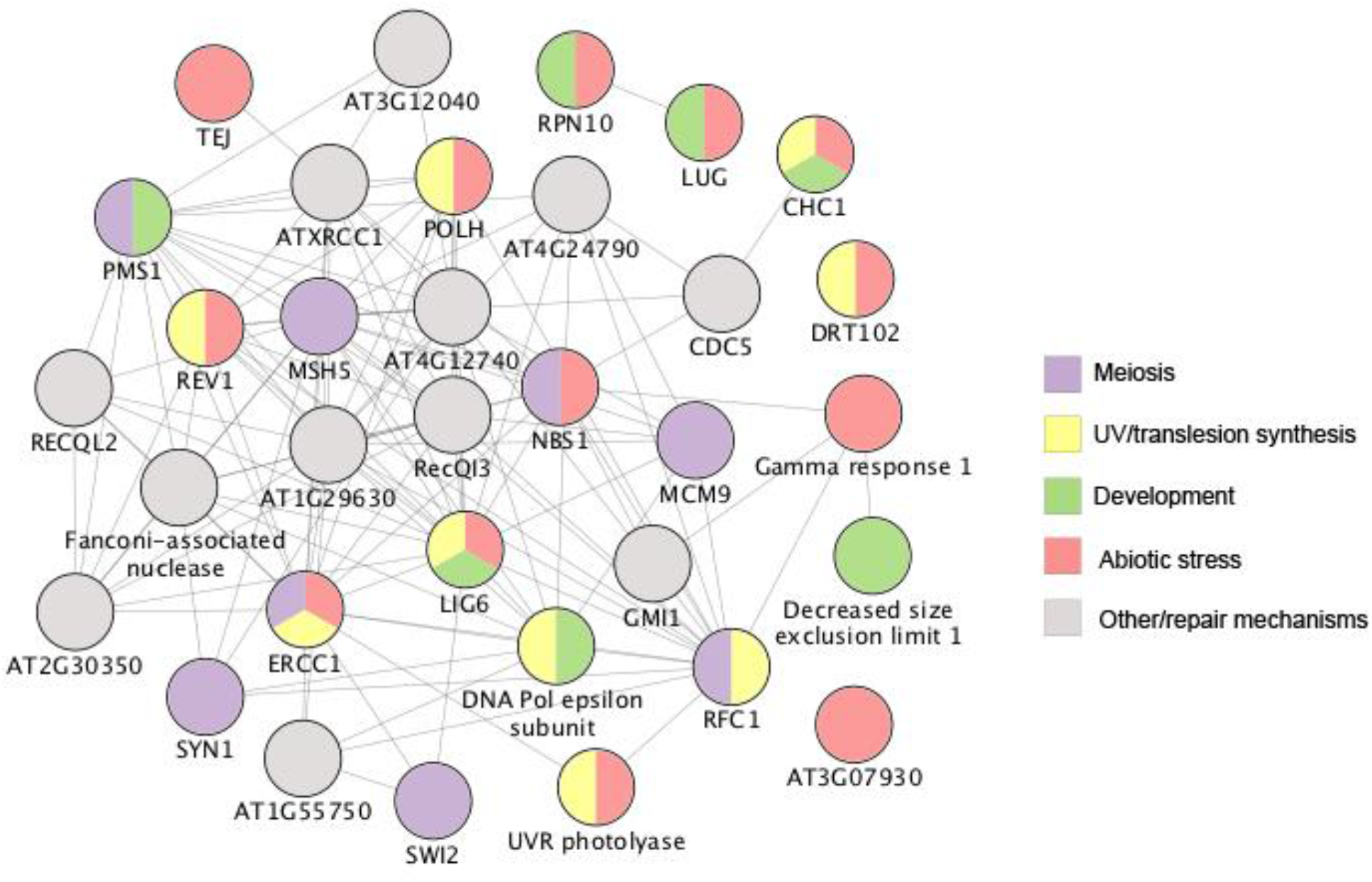
*A. negundo* gene families with ontology related to DNA damage and repair, and secondary enrichments categorized by color. Circles with multiple colors indicate multiple ontology assignments. Lines indicated known or predicted interactions, or other association via text-mining, co-expression, or protein homology. https://gitlab.com/PlantGenomicsLab/AcerGenomes/-/blob/master/acer/supplemental/figures/acne_dnadamage.jpg

In addition to UV-related DNA repair, there is another potential adaptation related to photooxidative stress. The early light induced protein 1 (ELIP) gene family stood out as significantly expanded in *A. negundo* relative to all other species, with twice as many genes compared to other *Acer*. Research has focused on ELIP’s protection against high light (Huang et al., 2012), but it also regulates seed germination in response to environmental factors (Rizza et al., 2011), and is also expressed in roots, and in response to some abiotic stressors including nitrogen in *Populus* (Luo et al., 2015), indicating additional protective functions. Capacity for regulation of photosynthesis and DNA repair mechanisms, within a largely streamlined genome, could represent investments with large-scale effects, altering growth according to resources or stress as described for *A. negundo* in various nutrient and light conditions.

## Conclusion

In this study we present new chromosomal-length genomes for *A. saccharum* and *A. negundo* and demonstrate the greater repeat, paralog, GC content, and expanding character of *A. saccharum* versus *A. negundo* which was reduced and rapidly contracting. We conducted a stem tissue expression study of *A. saccharum* subjected to long-term aluminum and calcium treatments. Larger trends related to the abiotic stress response and calcium deficiency were revealed against a complex background of expression. Gene families related to aluminum stress resistance and tolerance were compared showing moderate expansion of families in *A. saccharum*. Gene family dynamics combined with physiological and expression results, point toward potential aluminum resistance mechanisms in effect at the root. Broader gene family comparisons revealed expansion in traits associated with invasiveness and stress tolerance in *A. negundo*. Release of these genomes and a complementary expression analysis of trees in the HBEF NuPert study shed further light on mechanisms of tolerance and potential adaptations of increasing importance due to climate change.

## Methods

### Sequencing

Leaf material for *A. saccharum* was collected from the University of Maryland College Park Campus (GPS location 38°59’16.0”N 76°56’32.8”W). *A. negundo* was sourced from the ForestGEO Forest Dynamics plot (tag # 91603) located in Smithsonian Environmental Research Center, Edgewater, MD. Leaves were dark treated for at least one day prior to collection and shipped to Amplicon Express Inc. (Pullman, WA, USA) for DNA extraction. Both species were sequenced with Pacific BioSciences Sequel v2.1 using a total of fourteen SMRTcells, with the SMRTbell® Express Template Prep Kit v1.0, insert size 40 Kb, library size selection of 66 Kb for *A. saccharum* and 55 Kb for *A. negundo*. The resulting short read sequencing consisted of two lanes of Illumina HiSeq 2500 150PE with insert sizes ranging from 500 to 600 bp (Genomics Resource Center, University of Maryland School of Medicine Institute for Genome Sciences, Baltimore, MD, USA). For Hi-C, DNA leaf material was collected from the same individuals and extracted with the Phase Genomics Proximo Hi-C Plant kit (Phase Genomics, Seattle, WA, USA). Sequencing consisted of two lanes of mid output Illumina NextSeq 500 75PE with an average of 150 M reads per species (Center for Genome Innovation (CGI) at the University of Connecticut, Storrs, CT, USA). For the RNA-Seq used in annotation, leaf samples were collected from one individual per species at two years of age grown at the Michigan State University greenhouse. Sequencing consisted of Illumina HiSeq at 100PE.

Stem tissue from *A. saccharum* was sampled from nine trees across two seasons from the Nutrient Perturbation study plots at HBEF (North Woodstock, NH, USA). A single sapling (dbh < 21 cm) was selected from three plots in each of the three treatments (calcium, control, aluminum). A total of 18 libraries (nine in Oct 2017 (Fall); nine in May 2018 (Spring)) were sampled (Table S2). Tissue was sampled from five years of growth measured by spring wood internodes. Sections were frozen in liquid nitrogen in the field and RNA were extracted after grinding tissues in liquid nitrogen. Extractions were run on Agilent Bioanalyzer Tapestation for quantification and RNA integrity. Two samples were unsuccessful (one aluminum and one calcium) from the spring sampling. Libraries were prepared using Illumina TruSeq Stranded mRNA and sequenced with Illumina NextSeq 500, 150PE (CGI).

### Draft genome assembly

In advance of assembly, genome size was estimated with Jellyfish v2.2.6 (21-mers) and GenomeScope (Vurture et al., 2017) using the trimmed Illumina short-reads. The *A. saccharum* reads were test assembled using Illumina short-read, raw and corrected long-reads, and a hybrid of both. Draft assemblies were evaluated in relation to expected genome size, contiguity (N50 and number of contigs), conserved seed plant orthologs, and genomic/transcriptomic read alignments.

The best draft assemblies leveraged the deep PacBio sequencing (*A. saccharum* 103x, *A. negundo* 141x) and prioritized assembling repetitive regions of the genome and resolving the heterozygosity, found in both species. During this phase of the process, two draft assemblies of comparable results were used to investigate scaffolding potential. One was created with FALCON (pb-assembly v0.0.6) a *de novo*, diploid-aware assembler for PacBio (Chin et al., 2016), and the other was done using Canu (v1.6) error-corrected PacBio reads as input (Koren et al., 2017) to Flye v2.3.7, an efficient haploid assembler that leverages repeat graphs with read alignment techniques to resolve areas of repetition (Kolmogorov, 2019).

Alternate heterozygous contigs (haplotigs) were separated from the primary assemblies using Purge Haplotigs v1.0.4 (Roach et al., 2018). To determine coverage, PacBio reads were aligned to the primary assemblies with Minimap2 v2.15-r911-dirty (H. Li, 2018). RepeatModeler v1.0.8 was used to create the repeat libraries for this analysis.

### Hi-C scaffolding

Long-range scaffolding of the FALCON/Purge Haplotigs assembly with Hi-C reads followed processes recommended for the following suite of tools (*Genome Assembly Cookbook*, 2019). HiC reads were aligned with BWA mem -5SP and PCR duplicates removed with samblaster v0.1.24 (Faust & Hall, 2014). Scripts from the Juicer v1.5.6 (Durand, Shamim, et al., 2016) pipeline were modified to identify the Sau3AI restriction enzyme. The resulting Hi-C alignment file was provided to 3D-DNA v180419 (Dudchenko et al., 2017) for scaffolding, leveraging different parameters for each species according to the differing draft assembly characteristics. In particular, --diploid was added to *A. saccharum* to address remaining under-collapsed heterozygosity. JuiceBox was used to visualize Hi-C mapping against each scaffold created by the different parameter tests to visually detect which incorporated the majority of the contigs into the expected 13 pseudo-chromosomes(Dudchenko et al., 2018; Durand, Robinson, et al., 2016).

### Genome annotation

RepeatModeler v2.01 (Flynn et al., 2020) and RepeatMasker v4.0.6 (Smit et al., 2013) were used to softmask the assembly. Trimmed RNA reads were aligned to the assembly with Hisat2 v2.1.0 (Kim et al., 2015). GenomeThreader v1.7.1 (Gremme et al., 2005) was used to align protein sequences (derived from *de novo* transcriptome assembly; -gcmincoverage 80, -dpminexonlen 20 -startcodon -finalstopcodon -introncutout). Structural gene prediction was executed with BRAKER2 v2.0.5 (Hoff et al., 2019). The process converted RNA-Seq alignments to exon support in GeneMark-ET v4.32 (Lomsadze et al., 2014), and combined this output with protein alignments for two rounds of training with AUGUSTUS v3.2.3 (Camacho et al., 2009; Stanke et al., 2006, 2008) to predict coding regions in the genome. Extensive filtering was performed on the predicted gene space using gFACS v1.1 (Caballero & Wegrzyn, 2019). Evaluation of structural annotations were conducted with BUSCO and the PLAZA CoreGF rosids database v2.5 (Van Bel et al., 2019; Veeckman et al., 2016). Transcriptomic alignments were used to identify where they fully overlapped BRAKER-based models or provided additional support to those that did not pass previous filtering criteria. Transcriptome assemblies were conducted *de novo* with Trinity v2.20 (Grabherr et al., 2011). EnTAP v0.8.0 (Hart et al., 2020) was used to frame-select, functionally annotate, and identify potential contaminants for filtering, including bacteria, archaea, opisthokonta, rhizaria, and fungi. The resulting translated protein sequences were clustered with USEARCH v9.0.2132 (Edgar, 2010) at an alignment identity of 0.90. Transcriptomic alignments created with GMAP v2019-06-10 (Wu & Watanabe, 2005) and gFACs were compared against the genome annotation using GffCompare v0.11.5 (Pertea, 2018).

Functional annotation was performed using EnTAP v0.9.1, a pipeline that integrates both similarity search and other annotation sources including gene family (eggNOG), protein domains (Pfam), gene ontology, and pathway assignment (KEGG). The following public databases were included: NCBI RefSeq Complete, EMBL-EBI UniProt, and *Arabidopsis* (TAIR11).

### Comparative genomics

The translated gene space of 22 plant species were used for gene family analysis. *Acer* included *A. saccharum, A. negundo*, and *A. yangbiense*. The remaining species were selected from high quality public annotations. Fourteen broadleaf trees (*Betula pendula, Carica papaya, Citrus clementina* and *sinensis, Eucalyptus grandis, Juglans hindsii* and *regia, Pistacia vera, Populus trichocarpa, Prunus persica, Quercus lobata* and *robur*, and *Tectona grandis*), plus one woody angiosperm, *Vitis vinifera*, were included, along with one gymnosperm, *Ginkgo biloba. Oryza sativa* Kitaake was the representative monocot, along with *Amborella trichopoda* and *Nymphaea colorata*, representing other more basal lineages, and *Arabidopsis thaliana*, being the primary plant model system. OrthoFinder v2.3.7 (Emms & Kelly, 2018) was used to generate orthogroups. Resulting gene counts per orthogroup for *A. negundo, A. saccharum*, and the mean of the combined three *Acer* were each compared to the mean of other species to identify potentially expanded, contracted, missing, and novel gene families. The initial delineation of expansion and contraction was set at 2-fold above the standard deviation. Full absence was verified with alignment of the *Arabidopsis* protein sequence against the assembly. CAFE v5 (Mendes et al., 2020) was used to identify rapidly evolving gene families. Values from the newick species tree produced by OrthoFinder were multiplied by 100 to prevent issues with rounding in CAFE, and the tree was made ultrametric using OrthoFinder. The poisson root frequency distribution was run three times on gene families filtered by size to ensure convergence of a lambda value representing birth and death rate. The selected lambda value was then used to run the large family set. Functional enrichment of the resulting families was obtained from gProfiler v:e99_eg46_p14_f929183 (Raudvere et al., 2019) using the annotation of the representative (longest) gene when aligned to *Vitis vinifera*. This well annotated woody angiosperm represents an ideal source having no recent WGD (Tang et al., 2008).

### Whole genome duplication and synteny analysis

Characterization of putative paralogs, including whole genome duplication was done as previously described (Qiao et al., 2019) using DupGen_finder to separate whole genome, tandem, proximal, transposed, and dispersed duplicates. Categorization can be helpful in speculating on the origin of duplication such as transposable element activity, localized replication error, larger segmental translocations, or ploidy events. The whole genome duplicate frequency distribution was plotted by Ks value for analysis of peaks. Microsynteny of the small peak of recent supposed whole genome duplication seen in *A. saccharum* was further analyzed with MCScanX-based collinearity scripts (Nowell et al., 2018), as well as overall macrosynteny between the three *Acer*.

### Differential expression analysis

HBEF NuPert plots were used as a basis for this study as they were designed to reflect acidity and calcium levels over time as described by Berger et al. 2001. They consist of 12 *A. saccharum* dominant plots near reference watershed 6, with four receiving annual AlCl_2_ treatments 12 times from 1995 to 2015, and CaCl_2_ treatments were applied to 4 other plots for 4 years, followed by applications of slow-release wollastonite in 1999 and 2015 (Table S7). Three samples were collected from aluminum, calcium, and unamended control plots as described in the Sequencing section. Trimmed reads for each of the sixteen successful HBEF-sourced libraries were aligned to the *A. saccharum* reference genome with Hisat2 v2.1.0, and read counts were extracted with htseq-count v0.11.2. The R Bioconductor package, DESeq2 v1.26.0 (Love et al., 2014), was used for the expression analysis with the calcium as the control in pairwise comparisons of unamended to calcium and aluminum to unamended, representing increasing levels of aluminum, and then aluminum to calcium, contrasting the extremes. P-adjusted values greater than 0.1 were filtered. Pairwise comparisons, specific to each season (fall and spring) as well as combined resulted in a total of nine sets. Gene Ontology (GO) enrichment was conducted with g:Profiler (database version e99_eg46_p14_f929183) using alignments of differentially expressed protein models to both *Vitis vinifera* (Phytozome v12.1) and *Arabidopsis thaliana* (TAIR11) as baseline annotations.

*A. thaliana* was used as a representative model for pathway analysis in the Genemania application for Cytoscape v3.8.0. and *V. vinifera* (NCBI taxon ID:29760) (Smoot et al., 2011) was used similarly with STRINGDB v.11 in Cytoscape v2.7.1. Differentially expressed proteins for each pairwise comparison were used to visualize the fold-change values in context of the supported pathways. Networks were constructed with a confidence score of 0.4 and 20 maximal additional interactions and additional networks for protein models reported to be responsive to calcium deficiency and aluminum biotoxicity were imported and merged.

## Data Availability

Sequencing for *A. negundo*, along with the genome, annotations, and RNA-Seq are available in BioProject PRJNA750066. Corresponding data for *A. saccharum* is in BioProject PRJNA748028 with the exception of RNA-Seq used in annotation which is available in PRJNA413418. RNA-Seq used in the differential expression study are available in BioProject PRJNA751902. Full details on assembly, annotation, gene family analysis, and differential expression analysis can be found at https://gitlab.com/PlantGenomicsLab/AcerGenomes.

## Acknowledgements

NGS was funded by the National Science Foundation (NSF DEB-2029997; NSF EF-1638488). Hi-C library preparation and sequencing was funded by the Ronald Bamford Fund Endowment for Ecology and Evolutionary Biology to the Department of Ecology and Evolutionary Biology, University of Connecticut. Support for the HBEF RNA-Seq was provided by the University of Connecticut Center for Biological Risk.

The authors would like to thank the Institute for Systems Genomics (ISG) at UConn for sequencing support, the Microbial Analysis Resources and Services (MARS) within the Center for Open Research Resources & Equipment (COR^2^E) at UConn for RNA extraction support, the Computational Biology Core for HPC services, and the Hubbard Brook Experimental Forest for access to the NuPert plots. Thanks to Laura Figueroa Corona for assistance with the range map, and Sumaira Zaman, Alison Scott, and Nasim Rahmatpour for sharing computational approaches and scripts. Thank you to Dr. Yaowu Yuan for advice on both data analysis and the manuscript. The *A. negundo* individual sequenced is a part of the Smithsonian Environment Research Center Forest Dynamics Plot, which is part of the Smithsonian’s ForestGEO network. We thank ForestGEO and Director Stuart Davies for encouraging tree genome research in the network and for facilitating and funding (NSF DEB-1046113; NSF DEB-1545761) as well as previous working group discussions on the topic. The authors declare no conflict of interest.

## Author Contributions

SM, NS, JW, PS, and UU conceived and designed various aspects of the study; JW and NS coordinated and managed the study; SM, UU, NS, PS, AT performed the sampling and experiments; AT performed transcriptomic analysis; SM performed the assembly, annotation, and related comparative genomic and expression analysis under the advisement of JW; SM, JW wrote the manuscript; UU, NS, and AT contributed sections. All authors read and approved the final manuscript.

## Supplemental

**Figure S1**. Genome size estimation using k-mer distribution analysis https://gitlab.com/PlantGenomicsLab/AcerGenomes/-/blob/master/acer/supplemental/figures/Figure_1_SuppInfo.pdf

**Figure S2**. Hi-C plots https://gitlab.com/PlantGenomicsLab/AcerGenomes/-/blob/master/acer/supplemental/figures/Figure_2_SuppInfo.pdf

**Figure S3**. PCA plot https://gitlab.com/PlantGenomicsLab/AcerGenomes/-/blob/master/acer/supplemental/figures/Figure_3_SuppInfo.pdf

**Figure S4**. Syntenic comparisons between the three *Acer* genomes https://gitlab.com/PlantGenomicsLab/AcerGenomes/-/blob/master/acer/supplemental/figures/Figure_4_SuppInfo.pdf

**Table S1**. Illumina, PacBio, and Hi-C sequencing data summaries https://gitlab.com/PlantGenomicsLab/AcerGenomes/-/blob/master/acer/supplemental/tables/sequencing_tables.pdf

**Table S2**. HBEF table of trees https://gitlab.com/PlantGenomicsLab/AcerGenomes/-/blob/master/acer/supplemental/tables/hbef_trees.pdf

**Table S3**. HBEF GO enrichment https://gitlab.com/PlantGenomicsLab/AcerGenomes/-/blob/master/acer/supplemental/tables/hbef_functional_enrichment.pdf

**Table S4**. Orthogroup statistics by species https://gitlab.com/PlantGenomicsLab/AcerGenomes/-/blob/master/acer/supplemental/tables/Orthogroup_summary_table.pdf

**Table S5**. OF dynamics GO enrichment https://gitlab.com/PlantGenomicsLab/AcerGenomes/-/blob/master/acer/supplemental/tables/orthogroup_functional_enrichment.pdf

**Table S6**. CAFE GO enrichment https://gitlab.com/PlantGenomicsLab/AcerGenomes/-/blob/master/acer/supplemental/tables/cafe_functional_enrichment.pdf

**Table S7**. HBEF Nutrient Perturbation Treatment Schedule https://gitlab.com/PlantGenomicsLab/AcerGenomes/-/blob/master/acer/supplemental/tables/HBEFNuPertTreatmentTable2015.pdf

**File S1**. Assembly output stats https://gitlab.com/PlantGenomicsLab/AcerGenomes/-/blob/master/acer/supplemental/files/assembly_statistics.xlsx

**File S2**. Annotation details https://gitlab.com/PlantGenomicsLab/AcerGenomes/-/blob/master/acer/supplemental/files/annotation_statistics.xlsx

**File S3**. Collinearity analysis of recent Ks peak of WGD frequency https://gitlab.com/PlantGenomicsLab/AcerGenomes/-/blob/master/acer/supplemental/files/Acer_microsynteny.xlsx

**File S4**. HBEF differentially expressed genes https://gitlab.com/PlantGenomicsLab/AcerGenomes/-/blob/master/acer/supplemental/files/HBEF_DEGs.xlsx

**File S5**. Orthofinder significant dynamics https://gitlab.com/PlantGenomicsLab/AcerGenomes/-/blob/master/acer/supplemental/files/orthofinder_dynamics.xlsx

**File S6**. CAFE significant rapid evolution https://gitlab.com/PlantGenomicsLab/AcerGenomes/-/blob/master/acer/supplemental/files/cafe_rapidly_evolving.xlsx

**File S7**. *A. negundo* vs *Acer* contracted or missing using longest overall gene as annotation https://gitlab.com/PlantGenomicsLab/AcerGenomes/-/blob/master/acer/supplemental/files/acne_contracedmissing_longestoverall.xlsx

**File S8**. *A. negundo* vs *Acer* contracted or missing using longest *Acer* gene as annotation https://gitlab.com/PlantGenomicsLab/AcerGenomes/-/blob/master/acer/supplemental/files/acne_contractedmissing_longestacer.xlsx

**File S9**. *A. negundo* vs *Acer* expanded or novel using longest overall gene as annotation https://gitlab.com/PlantGenomicsLab/AcerGenomes/-/blob/master/acer/supplemental/files/acne_expandednovel_longestacer.xlsx

**File S10**. *A. negundo* vs *Acer* expanded or novel using longest *Acer* gene as annotation https://gitlab.com/PlantGenomicsLab/AcerGenomes/-/blob/master/acer/supplemental/files/acne_expandednovel_longestoverall.xlsx

**File S11**. *A. saccharum* vs *Acer* contracted or missing using longest *Acer* gene as annotation https://gitlab.com/PlantGenomicsLab/AcerGenomes/-/blob/master/acer/supplemental/files/acsa_contractedmissing_longestacer.xlsx

**File S12**. *A. saccharum* vs *Acer* contracted or missing using longest overall gene as annotation https://gitlab.com/PlantGenomicsLab/AcerGenomes/-/blob/master/acer/supplemental/files/acsa_contractedmissing_longestoverall.xlsx

**File S13**. *A. saccharum* vs *Acer* expanded or novel using longest *Acer* gene as annotation https://gitlab.com/PlantGenomicsLab/AcerGenomes/-/blob/master/acer/supplemental/files/acsa_expandednovel_longestacer.xlsx

**File S14**. *A. negundo* verified missing orthogroups https://gitlab.com/PlantGenomicsLab/AcerGenomes/-/blob/master/acer/supplemental/files/acne_missing_verified_string_mapping.tsv

**File S15**. *A. saccharum* verified missing orthogroups https://gitlab.com/PlantGenomicsLab/AcerGenomes/-/blob/master/acer/supplemental/files/acsa_missing_verified_string_mapping.tsv

**File S16**. Orthogroup comparisons for HBEF DEG and Al tolerance genes https://gitlab.com/PlantGenomicsLab/AcerGenomes/-/blob/master/acer/supplemental/files/hbef_orthogroup_Al_comparisons.xlsx

